# LuxHS: DNA methylation analysis with spatially varying correlation structure

**DOI:** 10.1101/2020.01.21.913640

**Authors:** Viivi Halla-aho, Harri Lähdesmäki

**Affiliations:** Department of Computer Science, Aalto University, FI-00076 Aalto, Finland

**Keywords:** DNA methylation, Bayesian analysis, Spatial correlation

## Abstract

Bisulfite sequencing (BS-seq) is a popular method for measuring DNA methylation in basepair-resolution. Many BS-seq data analysis tools utilize the assumption of spatial correlation among the neighboring cytosines’ methylation states. While being a fair assumption, most existing methods leave out the possibility of deviation from the spatial correlation pattern. Our approach builds on a method which combines a generalized linear mixed model (GLMM) with a likelihood that is specific for BS-seq data and that incorporates a spatial correlation for methylation levels. We propose a novel technique using a sparsity promoting prior to enable cytosines deviating from the spatial correlation pattern. The method is tested with both simulated and real BS-seq data and compared to other differential methylation analysis tools.

## 1 Introduction

DNA methylation is an epigenetic modification of the DNA where a methyl group is attached to a cytosine of the DNA. This phenomenon is essential for normal function of eukaryotic cells, and abnormal DNA methylation levels have been linked to diseases and cancer. DNA methylation is known to be a spatially correlated phenomena. In some cases, however, one or more cytosines in a local neighbourhood can deviate from the spatial correlation pattern due to e.g. transcription factor binding [4].

Many of the tools for differential methylation analysis assume spatial correlation without allowing cytosines to deviate from a common spatial correlation pattern. This inflexibility can lead us to not detecting all the possibly differentially methylated cytosines and could muddle the evidence for the non-deviating cytosines as well. For example RADMeth [3], which uses beta-binomial regression and weighted Z test and M^3^D [8] where maximum mean discrepancies over the regions are used for p-value calculation do not support finding deviating cytosines. One of the tools that could take such deviation into account is BiSeq [7] which has a hierarchical procedure, where defined CpG clusters are first tested by taking the spatial correlation into account and then trimming the found differentially methylated regions (DMRs) by removing the not differentially methylated cytosines from the regions. Even though spatial correlation is assumed in the first testing phase and preprocessing of the data includes smoothing, the second step allows for controlling location-wise false discovery rate (FDR). Also, BiSeq tool divides a DMR into smaller regions if the sign of the methylation difference changes.

In [5] we proposed a novel method LuxUS, that assumes spatial correlation for cytosines in a genomic window of interest. However, the method does not support detecting deviating cytosines and it calculates one Bayes factor for the whole genomic window. Here we present a different formulation of the spatial correlation that enables the analysis of deviating cytosines by introducing weight variables *d*_*i*_ for each cytosine *i* in the genomic window. The weight variable will tell whether the corresponding cytosine follows the general spatial correlation pattern or not. Horseshoe priors [2] are often used to enhance sparsity of the coefficients in generalized linear models, where the number of covariates in the model is very high and it is assumed that many of them have little effect on the predicted variable. Here we utilize horseshoe prior in the definition of the weight variables *d*_*i*_. The statistical testing, i.e. calculation of Bayes factors, is done for each cytosine separately.

## 2 Methods

In this section the LuxHS model and analysis workflow is described. The model consists of a data generating process and a generalized linear mixed model, which models the methylation proportions by taking into account covariates, replicate effects and spatial correlation. After this the fitting of the model parameters and the method for testing for differential methylation is explained.

The first step of workflow in LuxHS analysis is to divide the data set of interest into genomic windows, which are then analysed one at a time with possibility of parallelisation. A simple preanalysis method for this purpose was proposed by Halla-aho and Lähdesmäki [5]. The cytosines are divided into genomic windows based on their genomic distance and a maximum number of cytosines in a window while filtering out cytosines with low coverage. Genomic windows with too low average coverage can also be filtered out. For windows with high enough coverage, an F-test is performed to quantify the significance for the variable of interest. The F-test p-value threshold is set to a moderate value to refrain from filtering out too many prospective genomic windows. The windows that passed the F-test phase are further processed into LuxHS input format. The model parameters are fitted, after which statistical testing is performed. Finally, the found differentially methylated cytosines can be combined into DMRs for which a follow-up analysis, such as gene-enrichment analysis, can be performed.

### 2.1 Model

Plate diagram of the model is presented in Fig. 1. The analysis of an experiment with *N*_*R*_ samples is performed for a genomic window with *N*_*C*_ cytosines at a time. The total sequencing read counts for each cytosine are stored in **N**_BS,tot_ out of which **N**_BS,C_ (both vectors of length *N*_*C*_ · *N*_*R*_) were methylated. If the experimental parameters for each sample are available, they can be stored in vectors **BS**_eff_, 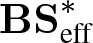 and **seq**_err_, each of length *N*_*R*_, which correspond to bisulfite conversion efficiency, incorrect bisulfite conversion efficiency and sequencing error. If the experimental parameters are not known, they can be set to correspond to a perfect experiment with no sequencing error and perfect bisulfite conversion efficiency.

**Figure 1:**
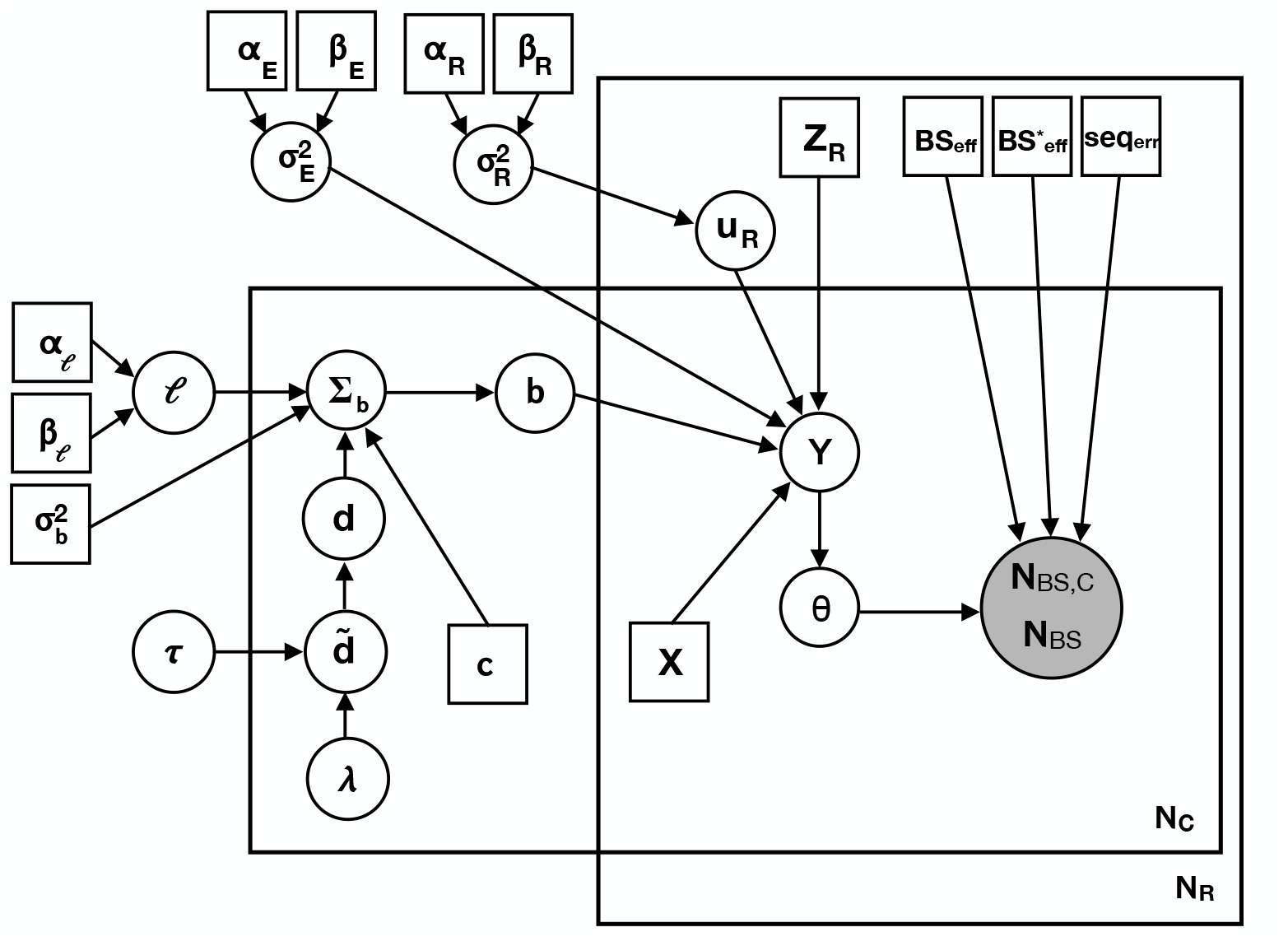
Plate diagram of the LuxHS model. Rectangles represent input data such as design matrices or fixed hyperparameters. White and grey circles represent latent and observed variables respectively.

The methylated cytosine count *N*_BS,C,*i*_ for observation *i*, *i* = 1, …, *N*_*R*_ · *N*_*C*_, follows binomial distribution

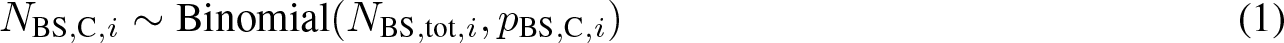

with success probability, e.g. probability of observing a C in bisulfite sequencing experiment, *p*_BS,C,*i*_, which is calculated as

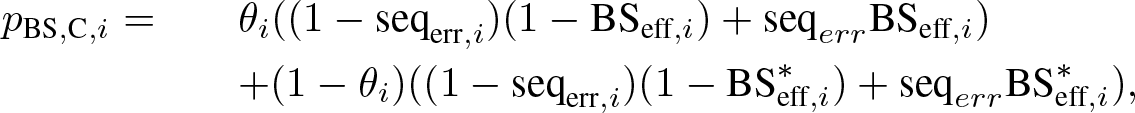

where *θ*_*i*_ is the methylation proportion for the observation *i*, *i* = 1, …, *N*_*C*_ · *N*_*R*_. seq_err,*i*_, BS_eff,*i*_ and 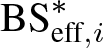 are the experimental parameters for the replicate corresponding to index *i*. The equation follows the probability tree in Fig. 2. This is the same data generating process as in LuxGLM [1] for non-methylated and methylated cytosines. Methylation proportions are estimated using the generalized linear mixed model of the form

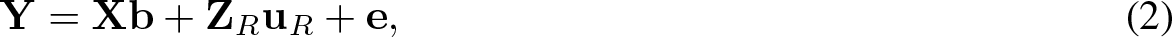

where term **Xb** is fixed effect, **Z**_*R*_**u**_*R*_ is replicate random effect and **e** is noise term with distribution 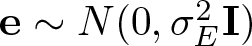 and prior 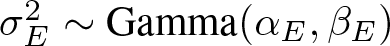. The number of covariates in the fixed effect term is *N*_*P*_. Each of the cytosines has its own set of fixed effect coefficient vector **b**_*j*_ of length *N*_*P*_, *j* = 1, …, *N*_*C*_, and thus 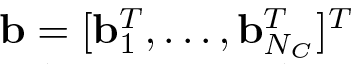 has length *N*_*C*_ · *N*_*P*_. The design matrix **X** size is (*N*_*C*_ · *N*_*R*_) × (*N*_*C*_ · *N*_*P*_) and it has the individual cytosine design matrices as block matrices in the diagonal. The fixed effect coefficients have prior distribution **b** ~ *N*(0, **Σ**_**b**_), where **Σ**_**b**_ is a covariance matrix. Matrix **Z**_*R*_ is the random effect design matrix of size (*N*_*C*_ · *N*_*R*_) × *N*_*R*_ and the vector **u**_*R*_ of length *N*_*R*_ contains the effects for each replicate. The effects have a normal prior distribution 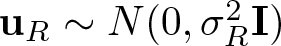, where 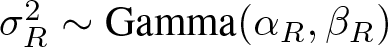 is the variance term for the replicate random effect.

**Figure 2:**
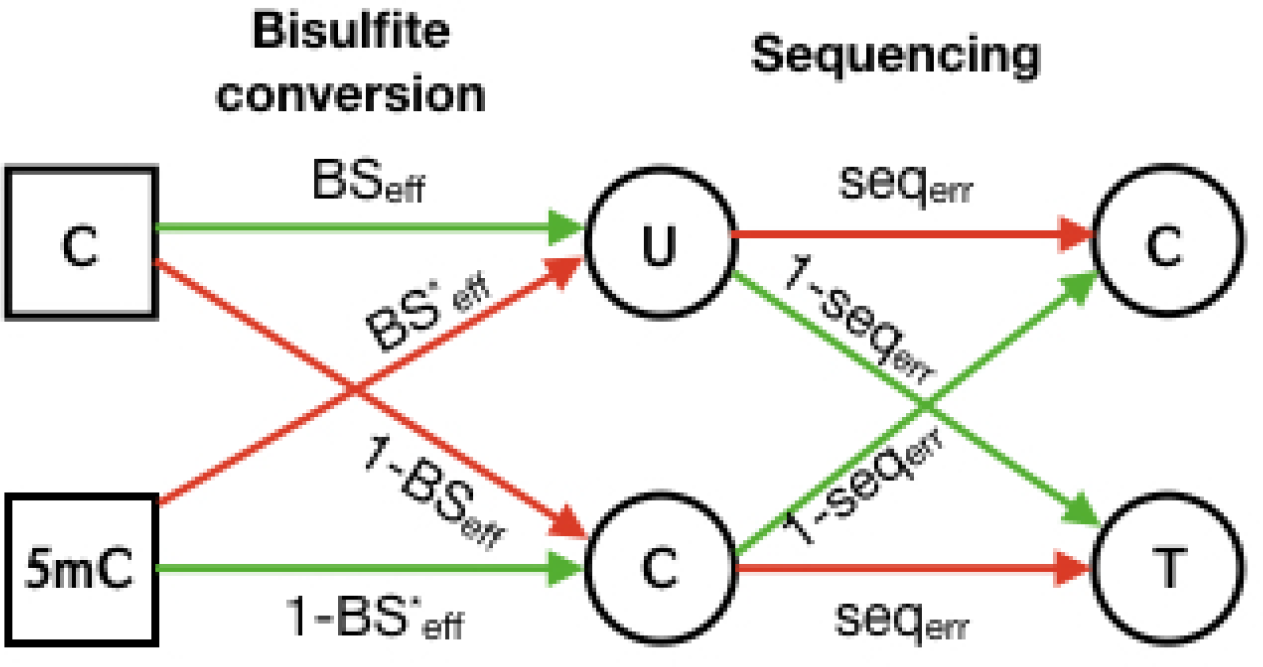
The probability tree for observing a C or T in bisulfite sequencing data when the true methylation state is methylated or unmethylated.

The spatial correlation structure is brought to the model through the fixed effect coefficients’ covariance matrix **Σ_b_**. Using indexing notation *b*_*j*, *k*_, *j* = 1, …, *N*_*C*_, *k* = 1, …, *N*_*P*_, to distinguish coefficients for each cytosine and covariate, **Σ_b_** can be expressed as

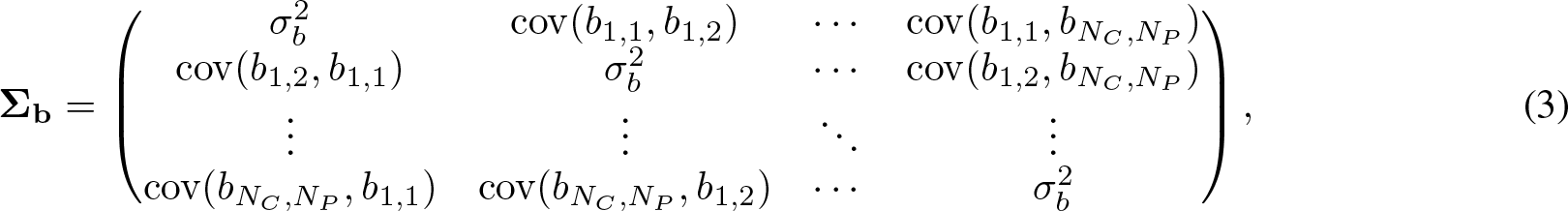

where the covariance terms are

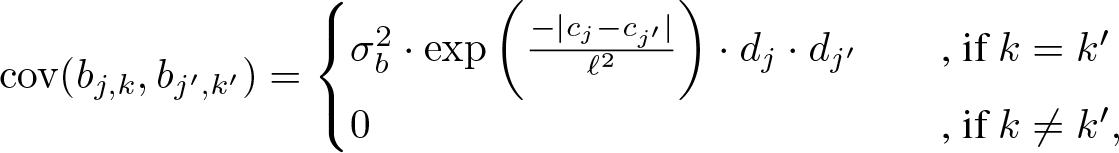

which gives the coefficients of different covariates zero covariance to fulfill linear model requirements. In the computation of the covariance terms, the cytosine locations *c*_*j*_ and *c*_*j′*_ and the lengthscale parameter *ℓ* with prior *ℓ* ~ Gamma(*α*_*ℓ*_, *β*_*ℓ*_) are used. The coefficient variance 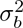 is set to the value 15. Weight variables *d*_*j*_ and *d*_*j′*_ tell whether the corresponding cytosine follows the correlation pattern along with its neighboring cytosines. The weight variables can have values ranging from 0 to 1, and they have the ability to scale down the covariance terms cov(*b*_*j*,*k*_, *b*_*j′*,*k′*_).

The correlation weight variable *d*_*j*_ for cytosine *j* = 1, …, *N*_*C*_ is calculated through transformation

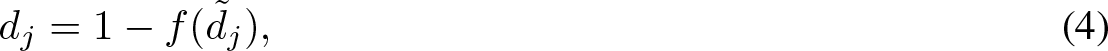

where transformation function *f* (*x*) is a generalized logistic function

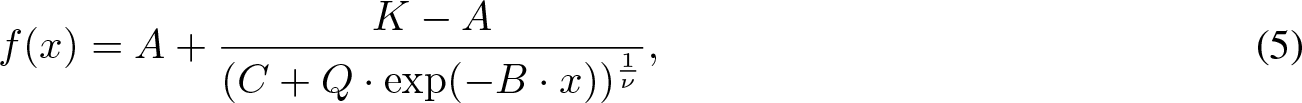

where *A* = 0, *K* = 1, *C* = 1, *Q* = 10, *B* = 5 and *ν* = 0.5. This transformation ensures that the resulting *d*_*j*_ have values from range [0, 1]. The auxiliary variable 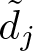 has a horseshoe prior with the modification of the normal priors for 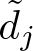 being restricted to the positive side, defined as

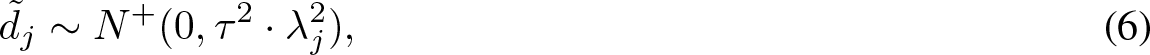

where the global shrinkage parameter *τ* and local shrinkage parameters λ*_j_* have positive Cauchy hyperpriors *τ* ~ *C*^+^(0, 1), and λ*_j_* ~ *C*^+^(0, 1). The level of sparsity of vector 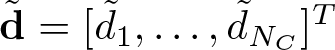 containing 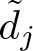, *j* = 1, …, *N*_*C*_, can be controlled with the choice of hyperprior for *τ*.

Finally, the methylation proportions *θ_i_* in Eq. 2 are calculated with the sigmoid function

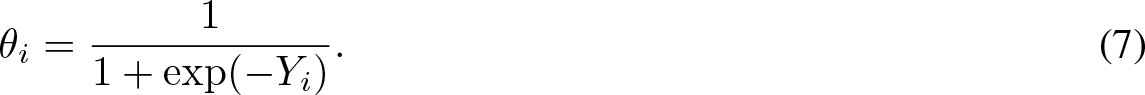

### 2.2 Fitting the model parameters with Stan and testing differential methylation

The model is implemented with probabilistic programming language Stan, and the Stan program is used for sampling from the posterior distribution. Stan offers both Hamiltonian Monte Carlo (HMC) and automatic differentation variational inference (ADVI) approaches for obtaining posterior samples and either one can be used for LuxHS. As variational inference approaches are often faster than Markov chain Monte Carlo (MCMC) methods such as HMC, they are a potential alternative to MCMC in computationally heavy tasks.

After obtaining samples for the model parameters, Bayes factors can be calculated for each cytosine to describe the evidence for two alternative models. The testing is done cytosine-wise, which enables deviating Bayes factor values inside a genomic window. There are two versions of the differential methylation test, with the type 1 test having a base model *M*_0_ : *b*_*j*,*k*_ = 0 and an alternative model *M*_1_ : *b*_*j*,*k*_ ≠ 0, subscript *j* corresponding to the cytosines *j* = 1, …, *N*_*C*_ and subscript *k* corresponding to the covariate of interest. The type 2 test has a base model *M*_0_ : *b*_*i*,*j*_ − *b*_*j*,*k′*_ = 0 and an alternative model *M*_1_ : *b*_*j*,*k*_ − *b*_*j*,*k′*_ ≠ 0, subscripts *k* and *k′* corresponding to the covariates of interest. Corresponding Bayes factors (BF) are used for the testing. As exact Bayes factors are intractable, Savage-Dickey estimates of the BFs are used instead. S-D estimate for the type 1 test is

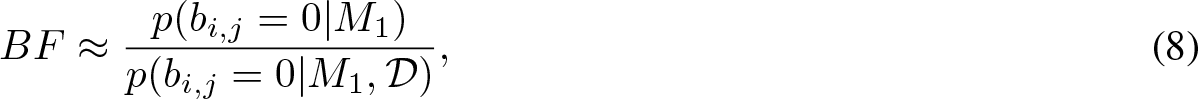

where 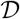 is the data. The numerator is calculated using the normal prior for **b** ~ *N* (**0**, Σ_****b****_) and the denominator is estimated from the obtained samples using kernel density estimation. The type 2 test S-D estimate is formed similarly.

## 3 Results

In this section we present the results for real and simulated BS-seq data sets. We first analyze whole genome bisulfite sequencing (WGBS-seq) data from [6] and demonstrate that LuxHS can identify differentially methylated cytosines as well as individual cytosines whose methylation state deviate from the general spatial correlation pattern. With simulated data (for which we know the ground truth) we quantitatively evaluate LuxHS performance and compare with other state-of-the-art methods.

### 3.1 Real bisulfite sequencing data

The colon cancer data set by Hansen [6] was used for testing LuxHS. The data set consists of six paired colon cancer and healthy colon tissue samples. The preanalysis step was run on data from chromosome 22 with the same settings as for LuxUS [5], and it resulted with 4728 genomic windows that passed the coverage and F-test criteria. Those genomic windows covered 86189 cytosines in total. For these windows, LuxHS analysis was performed. The LuxHS BF value distribution consisting of all 86189 cytosines is shown in Fig. 3. The histogram demonstrates that large majority of the Bayes factors had value smaller than 10 (or log(*BF*) ≤ 1), but there are also a few cytosines with high BF values indicating differential methylation. In total 5334 cytosines had BF> 3. To filter the results even further, a threshold for the minimum average difference between the case and control sample methylation states can be applied. In comparison, LuxUS analysis resulted in 593 windows (covering 10324 cytosines) with BF> 3, more detailed description of the results can be found from [5].

**Figure 3:**
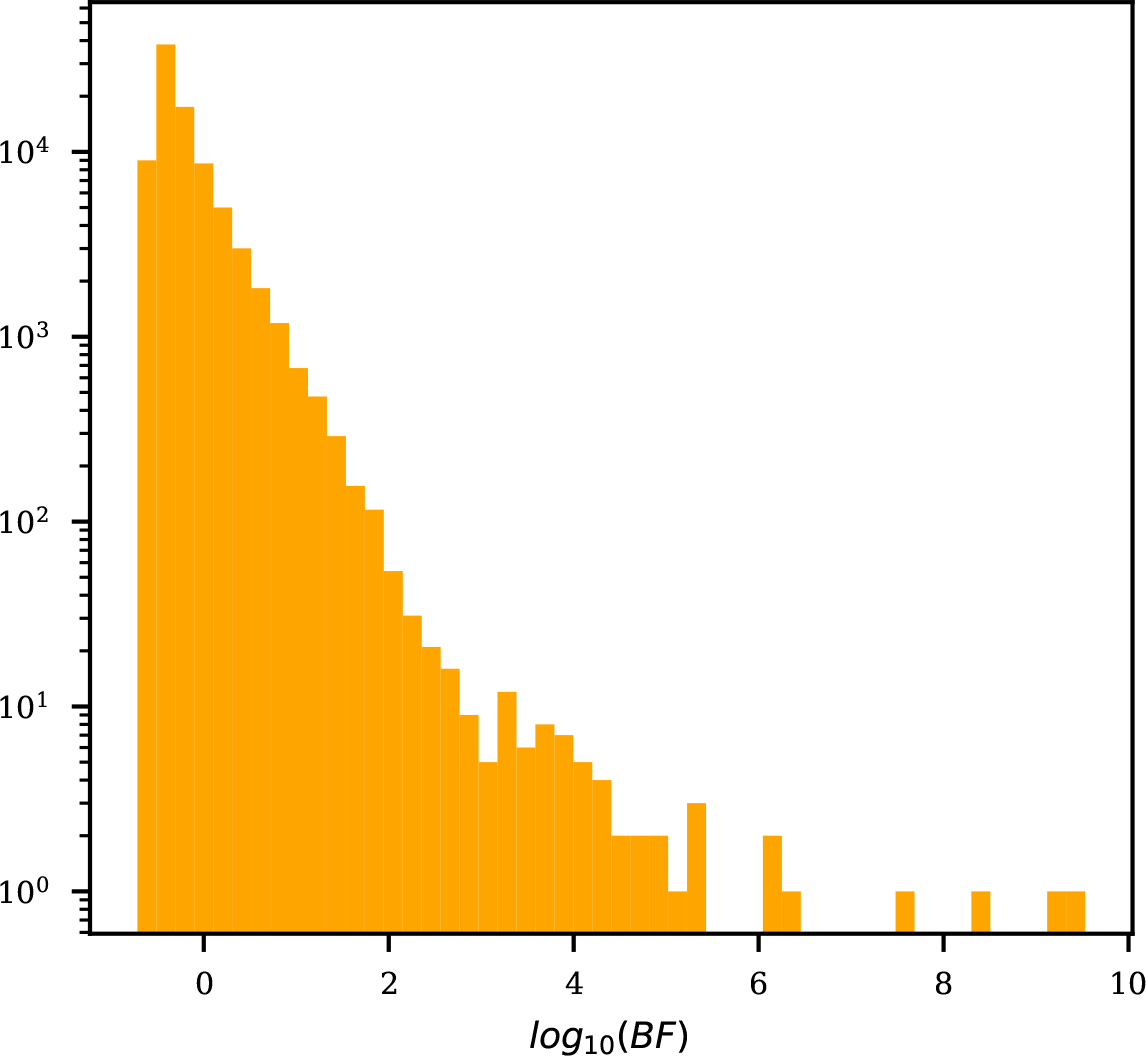
Histogram of the log(BF) values for the colon cancer data set. The y-axis of the histogram is in log-scale.

The number of cytosines for which the weight variables *d* were below 0.5 was 464. The Figure 4 demonstrates the differences between LuxUS and LuxHS results for a genomic window chr22:27014415-27015343. LuxUS gives one BF value (1.480) for the whole window, which suggests that there is no statistically significant differential methylation in the region. In contrary, LuxHS gives a Bayes factor for every cytosine separately while at the same time achieving two important goals: utilizing spatial correlation across the whole window of interest, and simultaneously detecting individual cytosines that deviate from this correlation pattern. Consequently, LuxHS is able to adapt to changes in the data swiftly. In the lowest panel of Fig. 4 it can be seen how LuxHS finds the cytosines for which the methylation states especially for the case samples are lower than in general in the window, and gives those cytosines lower weight parameter *d* values.

**Figure 4:**
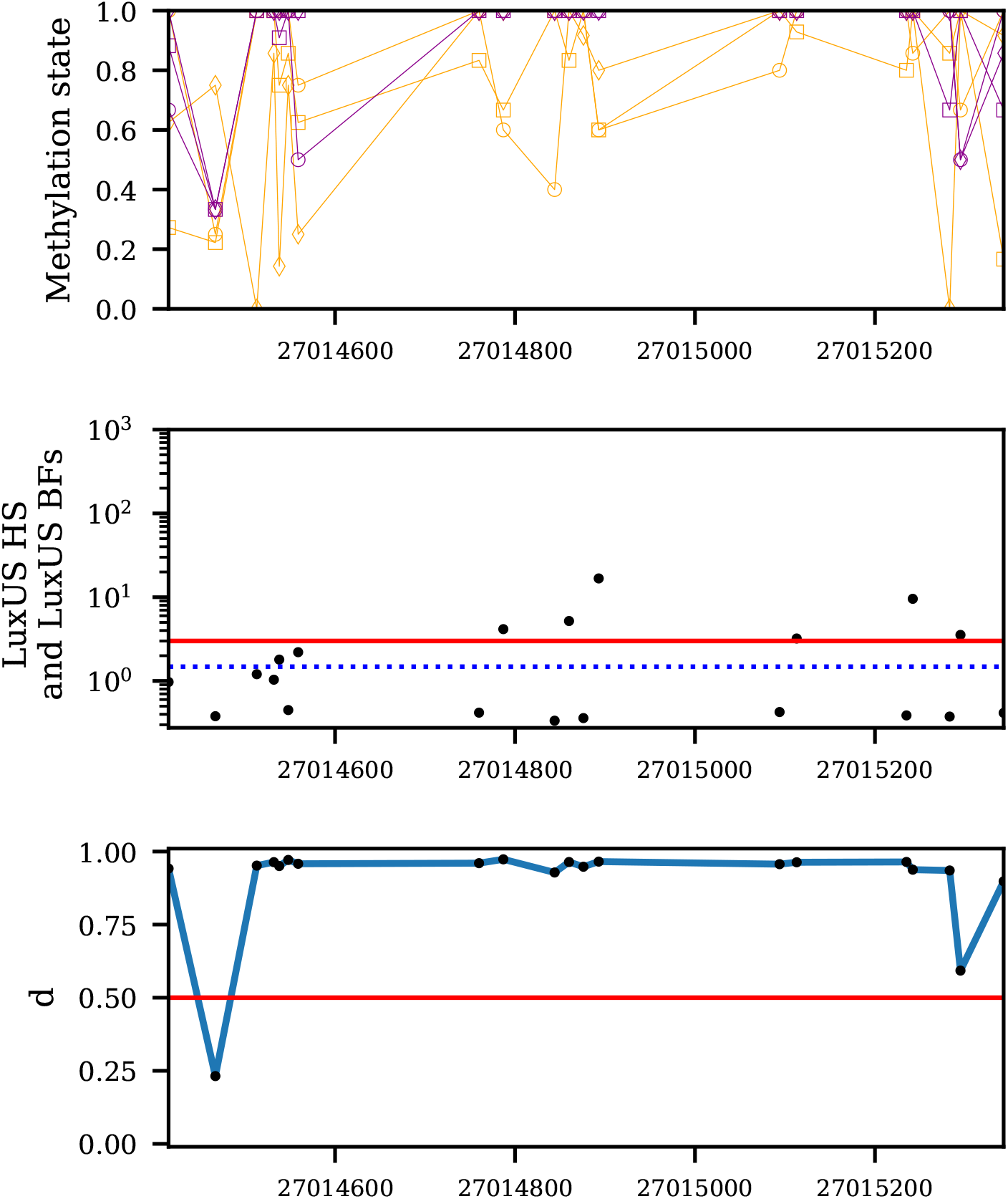
Results for a genomic region chr22:27014415-27015343. The top panel shows the methylation state data (as fractions *N*_BS,tot_/*N*_BS,C_) for the cytosines included in this genomic region. The cases have been plotted with purple and controls with orange, each replicate pair with a marker of its own. In the middle panel the LuxUS BF for the same region is plotted with the dashed blue line. The red line shows the threshold of BF value 3. The black dots are the LuxHS Bayes factors for each cytosine. The lowest panel shows the posterior mean of the samples for *d*, red line is plotted at value *d* = 0.5.

### 3.2 Simulated data

The data simulation was done using the LuxHS model, using variances 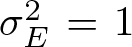, 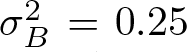 and 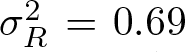. The experimental design of the simulations included an intercept term and a case-control binary covariate. The used coefficient mean *μ*_*B*_ values were [−1.4, 1], [−1.4, 1.8], [−1.4, 2.3] and [−1.4, 2.8], corresponding to methylation state differences between the case and control groups Δ*θ* values 0.2, 0.4, 0.5 and 0.6 respectively. The data is generated for type 1 tests. The number of total reads *N*_BS,tot_ for the methylated counts generation and the number of replicates *N*_*R*_ both had values 6, 12 and 24. For each combination of *μ_B_*, *N*_BS,tot_ and *N*_*R*_ we generated 100 data sets (each containing a genomic window of width 1000bp with 10 cytosines at randomly chosen locations) with and without differential methylation. The deviating cytosines had both opposite differential methylation status and deviating methylation state. We simulated data sets with 0, 1 and 2 deviating cytosines per data set. In this section we will refer to the number of deviating cytosines as *N*_*D*_.

LuxHS model was compared to four other models and tools: LuxUS [5], LuxUS applied separately to every cytosine (cytosine random effect removed from the model), RADMeth [3] and BiSeq [7]. Also, LuxHS models estimated with HMC and ADVI approaches are compared. BiSeq and RADMeth were ran with default settings. We decided not to present the BiSeq results, as BiSeq did not perform very well with the simulated data. This is perhaps due to the small size of the simulated genomic regions. The comparison was done with Receiver Operating Characteristic (ROC) curve statistics for all method. RADMeth runs resulted in a few NaN p-values, which were removed from the AUROC and TPR calculation. Area Under ROC curve (AUROC) value tables for *N*_*D*_ = 1 and *N*_*D*_ = 2 in Table 1-2 show that when the magnitude of differential methylation Δ*θ* is smaller, LuxUS performs the best. When Δ*θ* is higher, LuxHS and LuxUS for each cytosine separately have the highest AUROC values.

**Table 1:** AUROC values for the simulated data set with one deviating cytosine (*N*_*D*_ = 1) with best value for each simulation setting in bold.

| $\mu_B = [-1.4, 1]$ | | | | | | | $\mu_B = [-1.4, 2.3]$ | | | | |
| --- | --- | --- | --- | --- | --- | --- | --- | --- | --- | --- | --- |
| $N_R$ | $N_{BS,tot}$ | LuxHS<br>HMC | LuxHS<br>ADVI | LuxUS<br>sep | LuxUS | RADMeth<br>(NaN values) | LuxHS<br>HMC | LuxHS<br>ADVI | LuxUS<br>sep | LuxUS | RADMeth<br>(NaN values) |
| **6** | **6** | 0.528 | 0.532 | 0.488 | **0.587** | 0.565 (29) | 0.821 | 0.809 | 0.747 | **0.836** | 0.808 (46) |
| **12** | **6** | 0.622 | 0.62 | 0.592 | **0.654** | 0.648 (0) | **0.894** | 0.874 | 0.856 | 0.872 | 0.839 (31) |
| **24** | **6** | 0.682 | 0.674 | 0.666 | **0.732** | 0.676 (20) | **0.959** | 0.947 | 0.947 | 0.893 | 0.887 (30) |
| **6** | **12** | 0.539 | 0.543 | 0.522 | 0.565 | **0.569** (0) | 0.835 | 0.811 | 0.783 | **0.836** | 0.796 (10) |
| **12** | **12** | 0.612 | 0.601 | 0.588 | **0.66** | 0.614 (10) | **0.917** | 0.904 | 0.899 | 0.883 | 0.871 (10) |
| **24** | **12** | 0.708 | 0.704 | 0.699 | **0.714** | 0.698 (0) | **0.975** | 0.967 | 0.967 | 0.896 | 0.899 (40) |
| **6** | **24** | 0.59 | 0.584 | 0.569 | **0.618** | 0.59 (10) | **0.828** | 0.812 | 0.792 | 0.812 | 0.791 (10) |
| **12** | **24** | 0.664 | 0.656 | 0.641 | **0.688** | 0.651 (30) | **0.906** | 0.894 | 0.894 | 0.852 | 0.839 (10) |
| **24** | **24** | 0.75 | 0.747 | 0.74 | **0.767** | 0.727 (10) | **0.974** | 0.969 | 0.97 | 0.891 | 0.884 (30) |

**Table 2:** AUROC values for the simulated data set with two deviating cytosines (*N*_*D*_ = 2) with best value for each simulation setting in bold.

| $\mu_B = [-1.4, 1]$ | | | | | | | $\mu_B = [-1.4, 2.3]$ | | | | |
| --- | --- | --- | --- | --- | --- | --- | --- | --- | --- | --- | --- |
| $N_R$ | $N_{BS,tot}$ | LuxHS<br>HMC | LuxHS<br>ADVI | LuxUS<br>sep | LuxUS | RADMeth<br>(NaN values) | LuxHS<br>HMC | LuxHS<br>ADVI | LuxUS<br>sep | LuxUS | RADMeth<br>(NaN values) |
| **6** | **6** | 0.554 | 0.544 | 0.537 | **0.568** | **0.568** (36) | **0.759** | 0.73 | 0.742 | 0.712 | 0.685 (53) |
| **12** | **6** | 0.618 | 0.607 | 0.61 | **0.622** | 0.599 (20) | **0.858** | 0.823 | 0.845 | 0.75 | 0.742 (20) |
| **24** | **6** | **0.702** | 0.687 | 0.697 | 0.679 | 0.678 (30) | **0.952** | 0.934 | 0.95 | 0.782 | 0.797 (30) |
| **6** | **12** | 0.563 | 0.538 | 0.556 | **0.579** | 0.564 (1) | **0.796** | 0.769 | 0.779 | 0.722 | 0.721 (30) |
| **12** | **12** | 0.599 | 0.585 | 0.6 | **0.606** | 0.586 (20) | **0.897** | 0.88 | 0.896 | 0.757 | 0.751 (10) |
| **24** | **12** | 0.686 | 0.68 | **0.692** | 0.635 | 0.639 (20) | 0.956 | 0.945 | **0.958** | 0.783 | 0.802 (10) |
| **6** | **24** | 0.557 | 0.555 | 0.553 | **0.565** | 0.538 (30) | **0.835** | 0.808 | 0.825 | 0.738 | 0.729 (10) |
| **12** | **24** | 0.648 | 0.641 | **0.651** | 0.622 | 0.588 (30) | 0.905 | 0.889 | **0.906** | 0.75 | 0.774 (10) |
| **24** | **24** | 0.696 | 0.695 | **0.702** | 0.658 | 0.639 (30) | **0.965** | 0.957 | **0.965** | 0.785 | 0.787 (10) |

Based on the AUROC values, HMC version of LuxHS performs consistently slightly better than ADVI. The strength of ADVI is its computational efficiency. The mean runtime (over the 200 generated genomic windows) of the HMC version ranged from 40 to 971 seconds (for *μ_B_* = [−1.4, 1], *N*_BS,tot_ = [6, 12, 24] and *N*_*R*_ = [6, 12, 24]), while for ADVI the range is 5 – 47 seconds. The computations were run on a computation cluster.

The accuracy of estimating whether a cytosine is deviating or non-deviating from the common spatial correlation pattern was assessed using the estimated weight variable values *d*_*j*_. The posterior means of all weight variables were computed, and AUROC and TPRs were determined. The results in Table 3 show, that overall LuxHS can determine the deviance status accurately, but it seems not to be able to find all of the deviating cytosines. This indicates, that LuxHS rather gives too high *d* values than too low.

**Table 3:**
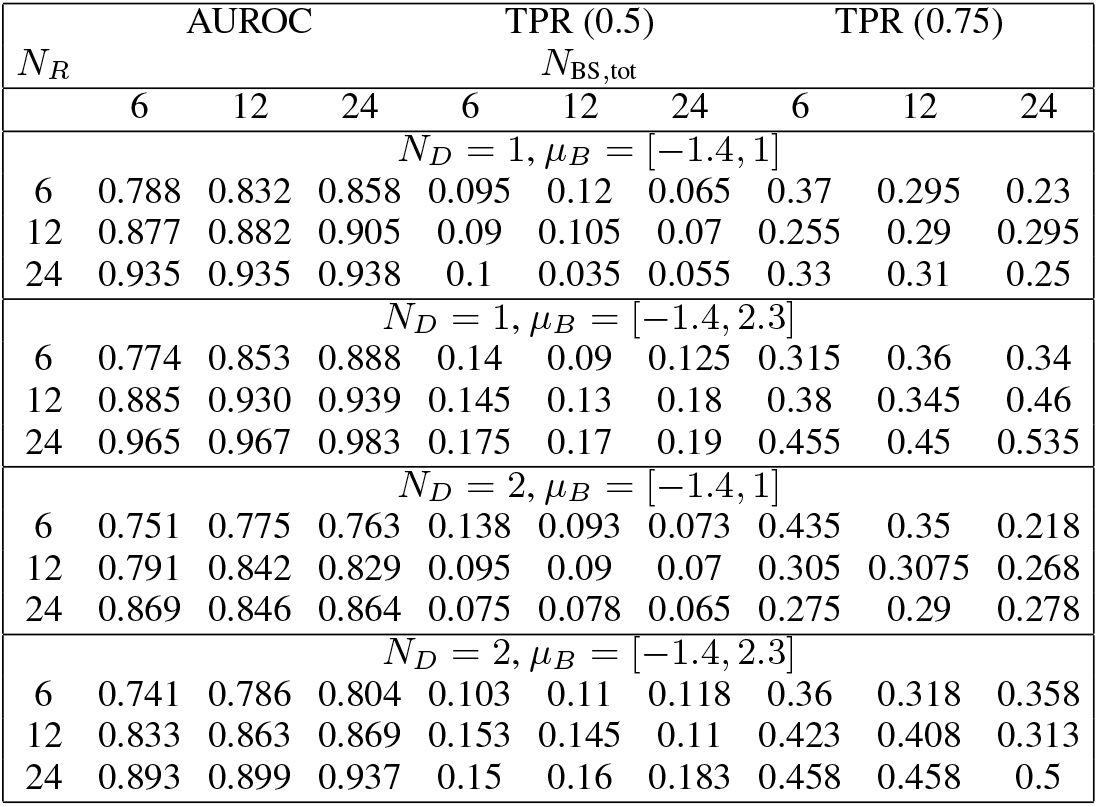
AUROC and true positive rates (TPR) for detecting the deviating cytosines in simulated data sets for LuxHS (HMC). AUROC was calculated using the posterior means for each *d*_*j*_. For the TPR calculation the *j*^th^ cytosine is considered deviating if the posterior mean of *d*_*j*_ is smaller than a threshold value. The results for two weight value thresholds 0.5 and 0.75 are shown in separate columns.

To investigate how LuxHS behaves when there are no deviating cytosines, such data sets were simulated and LuxHS analysis was performed along with the other methods it was compared to earlier. The data was simulated with Δ*θ* = 0.5. Out of the compared methods, LuxUS had the best AUROC values (see Table 4). LuxHS showed relatively good performance, demonstrating that the added flexibility of modeling cytosines that can deviate from the general spatial correlation pattern does not significantly decrease the performance of differential methylation analysis in the case of all cytosines following the same correlation pattern. Recall that for the cases where one or more of the cytosines deviate from the spatial correlation pattern LuxHS can reach state-of-the-art performance (see Tables 1-2). Moreover, LuxHS does not impose small values of weight variables *d*_*j*_ where it is not appropriate. There were no *d*_*j*_ values smaller than 0.5 for any of the generated genomic windows in any of the simulation settings.

**Table 4:**
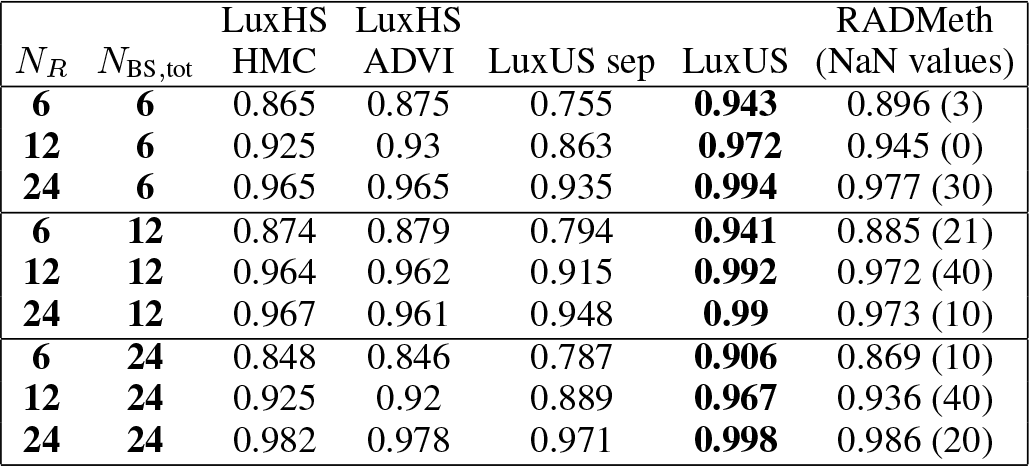
AUROC values for the simulated data set with zero deviating cytosines. *μ_B_* = [−1.4, 2.3] was used for the simulations. The best AUROC for each simulation setting is shown bolded.

## 4 Discussion

The analysis of real and simulated BS-seq data shows that LuxHS model can detect loci where the methylation state deviates from the surrounding cytosines. The tests with the simulated data show that the way LuxHS calculates Bayes factors separately for each cytosine can improve the accuracy when compared to LuxUS or other state-of-the-art methods, especially if the proportion of deviating cytosines is high.

The proportion of deviating cytosines that can be found in a genomic window could be further tweaked through the choice of hyperprior for global horseshoe prior *τ*. For example, the recommendations in [9] could be used if the default prior does not match the user’s beliefs about the number of deviating cytosines.

The covariance structure with possibility of breaking the correlation pattern might also be advantageous in other bioinformatic modeling purposes, where a spatial correlation pattern with possibility of deviation is needed. The spatial correlation structure proposed in here can be easily applied in a general or generalized linear model setting. Another application could be time series analysis, where consecutive time points are often correlated, but some of the time points may deviate from the expected correlation pattern e.g. due to an outlier value.

## 5 Conclusion

In this work we propose a novel method for differential methylation analysis, LuxHS. The tool supports detecting cytosines, which do not follow the same methylation pattern as its neighboring cytosines. This could happen because of e.g. transcription factor binding. The results with simulated and real BS-seq data show, that LuxHS is able to detect such cytosines and that this feature increases the accuracy of differential methylation analysis, especially when the number of deviating cytosines or the amount of differential methylation is higher. The tool and usage instructions are available in GitHub repository in https://github.com/hallav/LuxUS-HS.

## Funding

This work has been supported by the Academy of Finland (project numbers: 292660 and 314445).

## Acknowledgements

The calculations presented above were performed using computer resources within the Aalto University School of Science “Science-IT” project.

## Notes

https://github.com/hallav/LuxUS-HS

